# Functional effect of indole-3 carbinol in the viability and invasive properties of cultured cancer cells

**DOI:** 10.1101/2023.05.06.539692

**Authors:** Andrea S. Baez-Gonzalez, Jaime A. Carrazco-Carrillo, Gabriela Figueroa-Gonzalez, Laura Itzel Quintas-Granados, Teresita Padilla-Benavides, Octavio D. Reyes-Hernandez

## Abstract

Cancer treatment typically involves multiple strategies, such as surgery, radiotherapy, and chemotherapy, to remove tumors. However, chemotherapy often causes side effects, and there is a constant search for new drugs to alleviate them. Natural compounds are a promising alternative to this problem. Indole-3-carbinol (I3C) is a natural antioxidant agent that has been studied as a potential cancer treatment. I3C is an agonist of the aryl hydrocarbon receptor (AhR), a transcription factor that plays a role in the expression of genes related to development, immunity, circadian rhythm, and cancer.

In this study, we investigated the effect of I3C on cell viability, migration, invasion properties, as well as mitochondrial integrity in hepatoma, breast, and cervical cancer cell lines. We found that all tested cell lines showed impaired carcinogenic properties and alterations in mitochondrial membrane potential after treatment with I3C. These results support the potential use of I3C as a supplementary treatment for various types of cancer.

## INTRODUCTION

Cancer is a disease characterized by uncontrolled and autonomous growth of cells, which invades tissues locally and distantly via metastasis. According to data published by the World Health Organization in 2020, cancer is a leading cause of death worldwide, causing nearly 10 million deaths that year (1). The most common types of cancer include breast, liver, and cervical cancer, with lung, liver, colorectal, gastric, prostate, skin, and breast cancer having the highest incidence rates globally for both sexes. The predominant and most lethal types of cancer affecting men are prostate, lung, colon, liver, and stomach cancer, while breast, cervical, liver, colorectal, and ovarian cancer are the most harmful types for women. Breast cancer is the most common type of cancer in women, with over 2,260,000 new cases reported in 2020 (2). The activation of the human epidermal growth factor receptor 2 (hER2), estrogen and progesterone receptors, and mutations in the BRCA1/2 gene are some of the main factors contributing to breast cancer development. Cervical cancer, on the other hand, is mostly linked to papillomavirus (HPV) infection, which is a widespread sexually transmitted virus. Risk factors that contribute to breast and cervical cancer development include age, obesity, alcohol and tobacco use, family history, exposure to radiation, reproductive history, and postmenopausal hormone therapy. On the other hand, hepatocellular carcinoma (HCC) is the most common type of primary liver cancer that occurs in patients with chronic liver diseases.

Current therapeutic strategies for cancer include surgery, radiation therapy, and chemotherapeutics, with chemotherapy being the most common for patients in advanced stages. However, chemotherapy attacks non-tumor cells and generates additional side effects, such as neurotoxicity or nephrotoxicity, as in the case of cisplatin (3). Natural compounds, such as Indole-3-carbinol (I3C), found in cruciferous vegetables (4, 5), have potential anti-tumor properties and are structurally diverse that render less toxic side effects (6, 7). I3C has been proposed as a potential effective agent in the treatment of various types of cancer, including breast and prostate cancer (4, 8-30). It is an aryl hydrocarbon receptor (AhR) agonist and has been suggested to suppress the growth of breast and prostate cancer cell lines *in vitro*. I3C and its condensation products, such as 3,3’-diindolylmethane (DIM) and indole[3,2-b]carbazole (ICZ), are considered anti-carcinogenic compounds, and their biosynthesis is illustrated in **Figure 1A** (4, 12, 25, 31, 32). The mechanism of action of I3C is not completely understood, but it is proposed to have a role in preventing cancer development (5, 16). *In vitro* studies have suggested that I3C can suppress the growth of breast and prostate cancer cell lines (14, 22, 27). I3C is an aryl hydrocarbon receptor (AhR) agonist, which is a cytosolic ligand-dependent transcription factor that can bind to synthetic or natural xenobiotics. While most AhR agonist ligands are toxic, I3C and other naturally synthesized compounds have also been shown to activate this receptor (33-35). The AhR has a complex functional role within the cell, including cell cycle modulation associated with several processes in cell proliferation and differentiation, gene regulation, tumor development, and metastasis (36). Studies have shown that I3C can prevent cell communication junctions in rat primary hepatocytes and induce cell cycle arrest in breast cancer cell lines (20, 36). In cervical cancer, I3C has been shown to activate the AhR and decrease cell proliferation, possibly through *UBE2L3* mRNA induction that leads to the ubiquitination of the oncoprotein E7 from the human papillomavirus, the main causal agent of cervical cancer (37). Another AhR endogenous ligand, β-naphthoflavone (BNF or 5,6-benzoflavone; **Fig. 1B**), has also demonstrated anti-cancer properties against mammary carcinoma (38) and cervical cancer cells (39). The anti-tumorigenic function of AhR contrasts with the effect of stimulating malignant transformation of tissues that has also been reported in AhR (40, 41).

**Figure 1.**
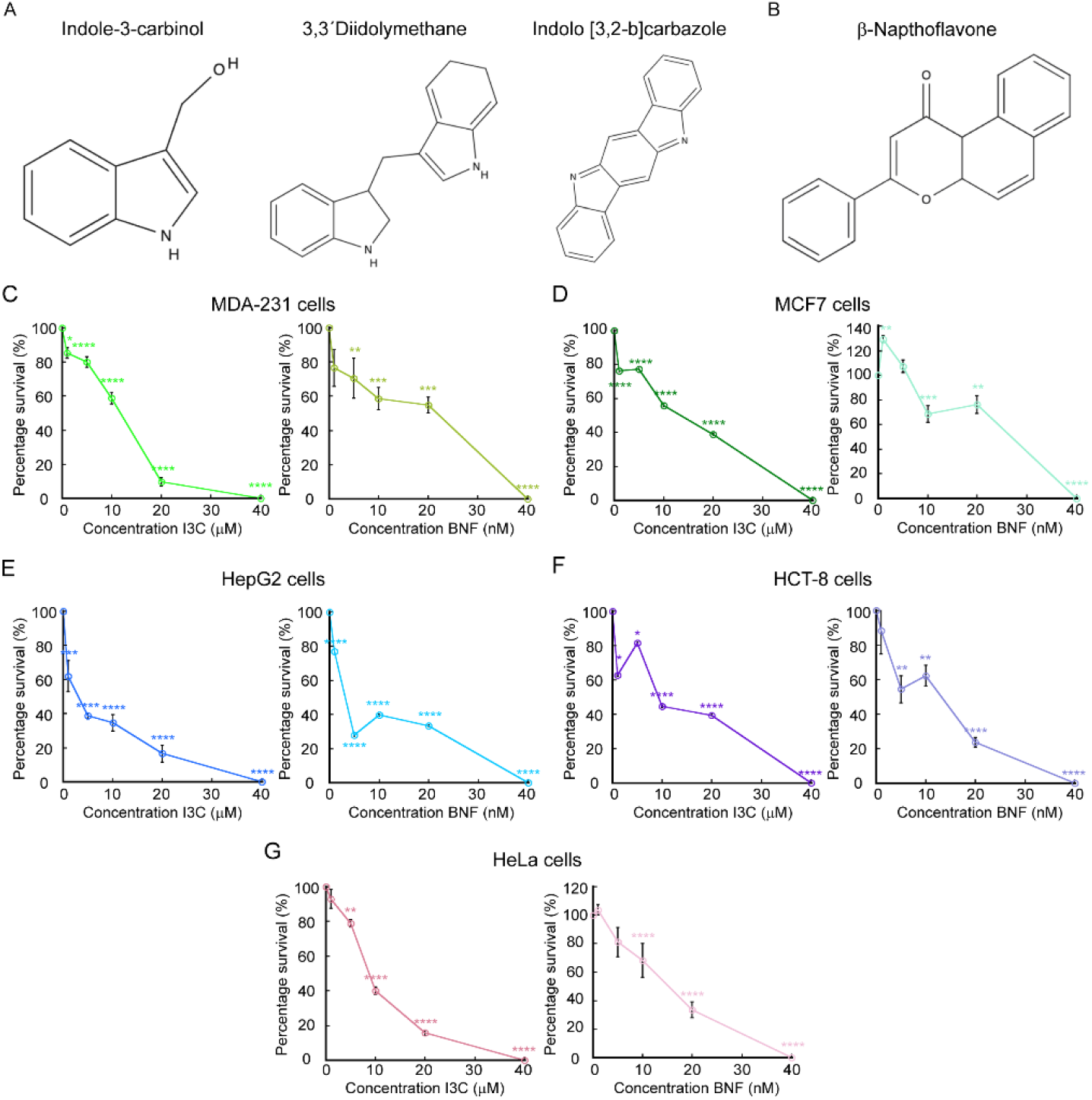
Indole-3-carbinol impairs proliferation of cancer cells. Chemical structure of Indole-3-carbinol (I3C) and two metabolite products (**A**) and β-Naphthoflavone (BNF, **B**). Cell counting assay of proliferating cancer cell lines grown for 48 h with increasing concentrations of I3c and BNF. Breast cancer cells MDA-MB-231 (**C**) and MCF-7 (**D**), the hepatocellular carcinoma cell lines HepG2 (**E**) and HCT-8 (**F**) and cervical cancer HeLa cells (**G**) were used. Data represent the percentage of survival of each strain under the presence of the two compounds. For all samples, data are the mean ± SE of three independent biological replicates. *p < 0.05; **p < 0.01; ***p < 0.001; ****p < 0.0001.

However, the effect of I3C on other types of cancer is not well understood. Thus, in this study, we aimed to evaluate the possible inducible anti-tumor potential of AhR through I3C in breast, cervical, and hepatocellular carcinoma cell lines. Each cell line presented different phenotypical characteristics that allowed us to investigate whether these compounds render similar positive effects against functional properties of different cancer models. We performed cytotoxicity assays to determine effective concentrations for I3C and functional assays such as wound healing and Matrigel invasion assays to determine the effect of I3C and BNF on the migratory and invasive capabilities of each cell line. Mitochondrial stress was also tested using the TMRE assays. Because of the effect exerted by I3C in the diverse carcinogenic hallmarks of the cells, we concluded that it has the potential to be used as supplemental natural therapy for the treatment of different types of cancer.

## MATERIALS AND METHODS

### Cell culture

The following cell lines were selected for this study: HeLa for cervical carcinoma model, HCT-8 and HepG2 for colon and hepatocellular carcinomas, MDA-MB-231 and MCF-7 cells for breast cancer model were obtained from American Type Culture Collection (ATCC, Manassas, VA, United States) and cultured in DMEM media (Sigma-Aldrich, St Louis, MO, United States) supplemented with 10% fetal bovine serum (FBS) and 1% antibiotics (penicillin G/Streptomycin, Gibco, Waltham, MA, United States) in a humidified atmosphere at 37º C and 5% CO_2_.

### Cell proliferation experiments

Cells were seeded on 24 well plates at 1×10^4^ cells/cm^2^ in DMEM supplemented with 10% FBS and antibiotics. Indole-3-carbinol (97%) and β-Naphthoflavone (98+%) were obtained from Thermo Scientific™ (122190250 and A18543.06, respectively). BNF was used as a reference compound of activation of AhR signaling. Both compounds were dissolved in DMSO and the I3C compound was used at 1, 5, 10, 20, 40 μM. BNF was tested at 1, 5, 10, 20, 40 nM concentrations. Cells were grown for 48 h in a humidified atmosphere at 37º C and 5% CO_2_. Then, the cells were washed gently with PBS, trypsinized and counted using a Cellometer Spectrum instrument (Nexcelom Biosciences). Four independent biological replicates were analyzed for each condition and cell line.

### Gene expression analyses

RNA was purified from three independent biological replicates of proliferating cancer cell lines with TRIzol (Invitrogen) according to the manufacturer’s instructions. cDNA synthesis was performed with 1 μg of RNA as template, using the HiFiScript gDNA Removal RT MasterMix (CoWin Biosciences) following the manufacturer’s protocol. Quantitative RT-PCR (qPCR) was performed with FastSYBR mixture (CoWin Biosciences) on the AriaMx Real-Time PCR System (Agilent Technologies) using the primers listed in **Table 1**. The delta threshold cycle value (ΔCT; (42)) calculated for each gene represents the difference between the CT value of the gene of interest and that of the control gene, *GAPDH*.

**Table 1.**
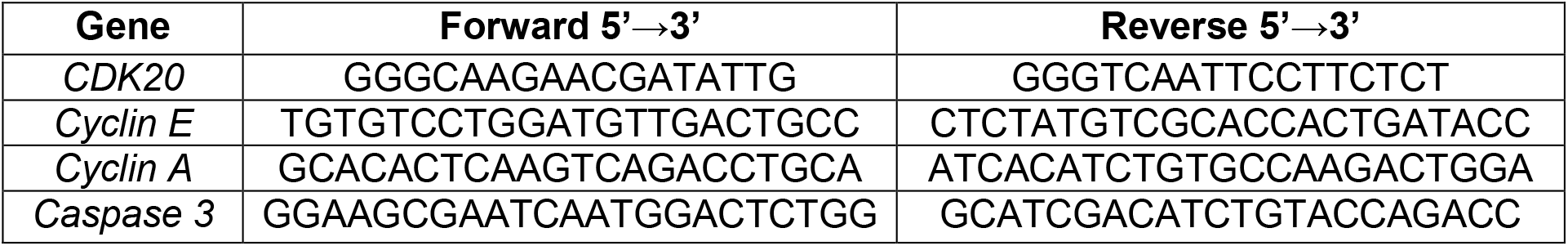
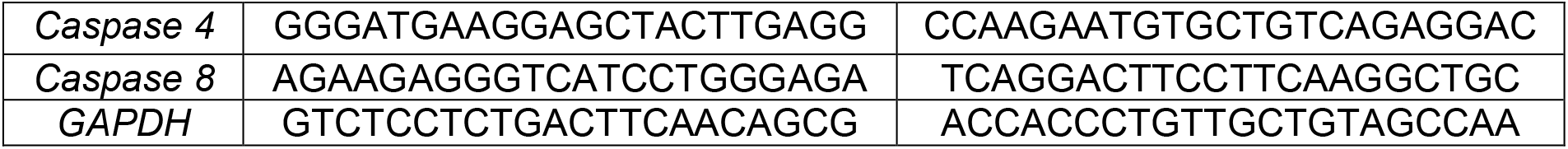
List of primers used for gene expression analyses.

### Wound Healing Assay

Cells were grown until confluence on 24 well plates in DMEM supplemented with 10% FBS and antibiotics. Cells were starved for 24 h in DMEM without FBS and cell proliferation was inhibited by treating the cells with 10 μM Cytosine β-D-Arabinofuranoside (AraC, C1768, Sigma-Aldrich) for 2 h. By inhibiting cell growth, we can ensure that the effect observed is only due to the movement of existing cells and not to new cells generated from cell division. The monolayers were then scratch-wounded using a sterile 200 μl pipette tip and suspended cells were washed away with PBS. The progress of cell migration into the wound was monitored every 24 h until wound closure using the ×10 objective of an Echo Rebel Microscope as previously described (43-45). The bottom of the plate was marked for reference, and the same field of the monolayers was photographed immediately after performing the wound (time = 0 h) and at different time-points after performing the scratch, as indicated in the figures. Area migrated by the cells was quantified using FIJI software, version 1.44p (46).

### Matrigel Invasion Assay

Matrigel invasion assay was performed following the Transwell chamber method as described (45, 47). Briefly, BioCoat® Matrigel® Invasion Chambers with 8.0 μm PET membranes (354481, Corning) were used to seed cells that were the indicated cell lines previously treated for 2 h with 10 μM AraC to inhibit cell proliferation. The cells were plated at 1.25 × 10^5^ cells/ml in 2 ml of serum-free medium on the top chamber, as recommended by the manufacturer. The lower chamber of the Transwell contained 2.5 ml of advanced DMEM supplemented with 10% FCS. Cells were incubated with I3C and BNF at the concentrations indicated in the figures. HeLa, MCF-7 and MDA-213 cells were allowed to invade the lower chamber for 24 h, while HCT-8 and HepG2 were incubated for 48 h. Then, the non-invading cells and Matrigel on the upper surface of the Transwell membrane were gently removed with cotton swabs. Invading cells on the lower surface of the membrane were washed and fixed with methanol for 5 min and stained with 0.1% crystal violet diluted in PBS. Images from 10 fields of three independent biological replicates were taken in ×10 objective of an Echo Rebel Microscope, and used for cell quantification using FIJI software, version 1.44p (46).

### Mitochondrial Membrane Potential

Changes in mitochondrial membrane potential produced by I3C and BNF in the five cancer cell lines was determined with the tetramethylrhodamine ethyl ester (TMRE) Mitochondrial Membrane Potential Assay Kit (ab113852, Abcam) following the manufacturer’s protocol. Briefly, proliferating cells were supplemented with 200 nM TMRE and incubated in the dark for 45 min at 37°C. The cells were then trypsinized and washed three times with PBS. Fluorescence intensity of TMRE was measured using a Spectrum Cellometer (Nexcelom Biosciences, Lawrence, MA, United States) by setting the filter excitation at 502 nm and emission at 595 nm, as previously reported (44, 48, 49). Data was analyzed with FCS Express 7 (De Novo Software).

### Statistical analyses

Statistical analyses were performed using Kaleidagraph (Version 4.1, Synergy Software). Statistical significance was determined using t-test where p < 0.05 was considered to be statistically significant.

## RESULTS

### I3C has an inhibitory effect in the proliferation of diverse cancer cell lines

I3C can impair the growth and induce apoptosis in certain cancer cell lines (8, 14, 16, 18, 19, 30, 31, 50). Therefore, we wanted to test additional effect of these drugs in characteristic functional hallmark properties of cancer cells. We selected cervical, breast, hepatic and colon carcinoma cell lines (HELA, MCF-7, MDA-MB-231, HEPG2, and HCT-8 respectively). These cells present different phenotypical characteristics and pathological mechanisms that allow to determine whether I3C and BNF would have additional anti-carcinogenic effects in different models. For instance, HeLa cells express an overactive type of telomerase, which prevents telomere shortening, prevents cell aging and death. The MCF-7 is an un-invasive cell line that express functional estrogen and epidermal growth factor (EGF) receptors, which makes it dependent on estrogen and EGF for growth. MDA-MB-231 cells are a model for more aggressive invasive, hormone-independent breast cancer, as it is triple negative for hormone receptors. HepG2 cells are non-tumorigenic cells with high proliferation rates and epithelial-like morphology that recapitulate hepatic functions *in vitro*. Insulin and the insulin-like growth factor II are typical markers of these cells. HCT-8 cells were isolated from the large intestine adenocarcinoma patient, these are adherent and present epithelial morphology. Both, HepG2 and HCT-8 present slower rates of migration and invasive capabilities.

First, we tested the cytotoxic effect of both compounds at increasing concentrations to determine the best working condition. To this end, each cell line was grown for 48 h under the presence of increasing concentrations of each I3C and BNF, as indicated in **figure 1C-G**. Supplementation of the culture media with I3C resulted in a decrease in proliferation of HeLa, MCF7, MDA-MB-231 and HCT-8 cell lines at concentrations of 10 μM. The hepatocellular carcinoma cell line HepG2 was sensitive to this compound at 5 μM of I3C. All cancer cell lines tested showed a high cytotoxic effect at a 40 μM concentration of I3C. In the case of BNF, we detected that cancer cell lines were sensitive to this compound at concentrations as low as 10 nM, with a larger effect in cell viability at 40 nM. Consistent with published data, our results suggested that both compounds have an inhibitory effect on the growth of different cancer cell lines. BNF seems to be more effective as the cells are sensitive to this compound at nM concentrations, while I3C exerts its effect at μM concentrations.

To investigate the mechanisms by which I3C impairs cell proliferation, we cultured cells with 10 μM and 10 nM of I3C and BNF for 48 h and analyzed changes in gene expression by qPCR. **Figure 2** illustrates that the cell lines responded differently to the compounds. We evaluated genes related to cell cycle regulation and found minimal changes under the conditions tested. For example, all cell lines showed decreased expression of *CDK20* when treated with BNF (**Fig. 2A-E**), and only the HCT-8 cell line exhibited decreased expression of this gene when treated with I3C (**Fig. 2D**). Cyclin E displayed a small but significant reduction in expression in HepG2 and HeLa cells (**Fig. 2C,E**), while Cyclin A was only reduced in HepG2 cells.

**Figure 2.**
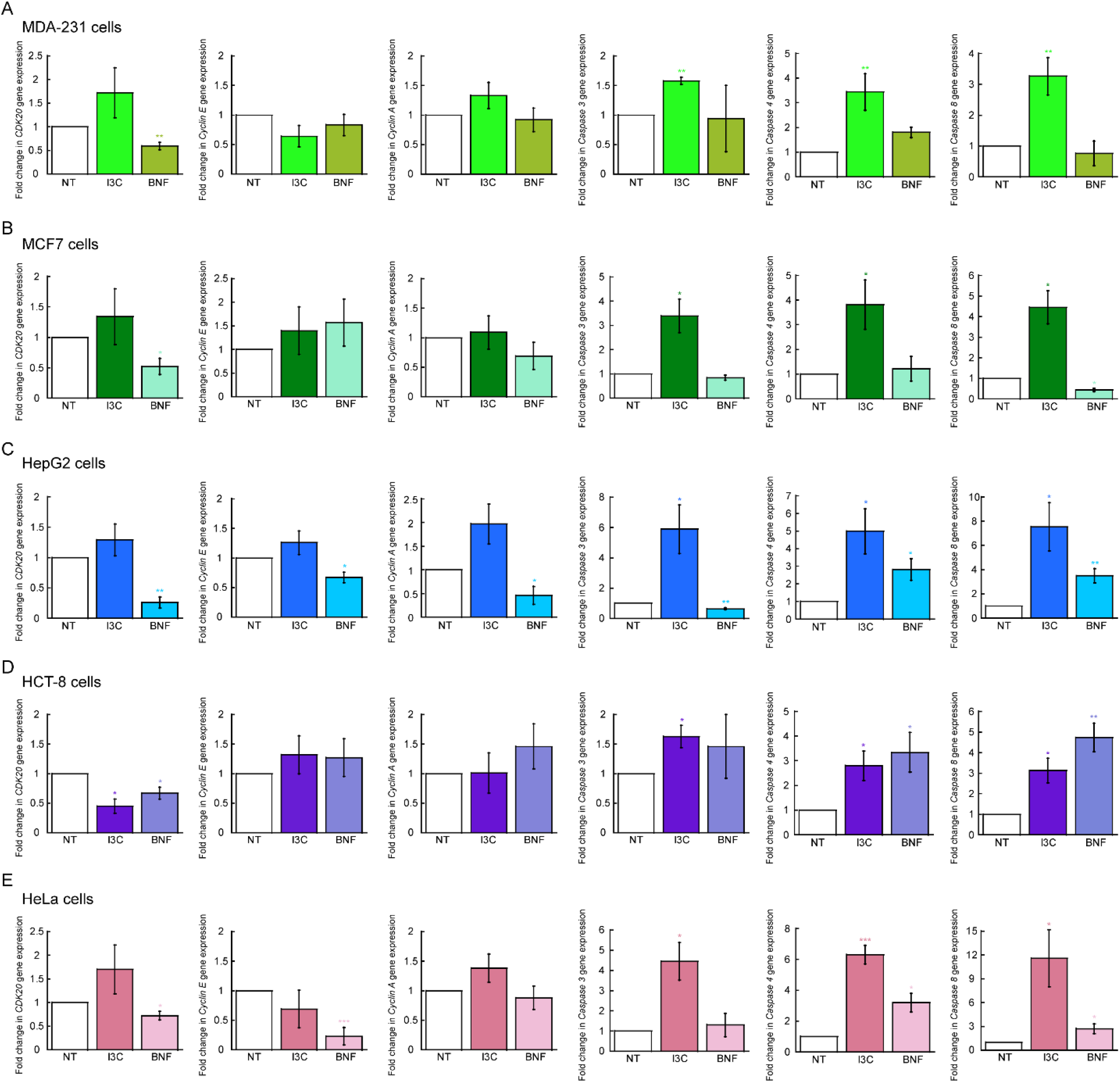
Effect of Indole-3-carbinole in the expression of cell cycle and apoptotic genes. Steady state mRNA levels determined by qRT-PCR of representative genes associated to cell cycle progression (*CDK20, Cyclin E*, and *Cyclin A*) and apoptosis (*Caspase 3, Caspase 4* and *Caspase 8*). Data represents the mean ± SE of three independent biological replicates. *p < 0.05; **p < 0.01; ***p < 0.001.

In contrast, apoptotic markers showed more drastic changes in cells treated with either compound. Caspase 3, 4, and 8 were significantly overexpressed in all cell lines when cultured with 10 μM of I3C. Interestingly, the expression of Caspase 4 and 8 was only elevated in HepG2, HCT-8, and HeLa cells upon BNF treatment. The data suggests that both compounds may mediate cell cycle progression and apoptosis through different mechanisms, potentially relying on specific cellular features. However, our results indicate that I3C primarily activates the apoptotic pathway, rather than mediating cell cycle progression. To further investigate the potential effects of these compounds on cell cycle regulation, we subjected the cells to mitotic arrest with Nocodazole, released them after 16 h, and monitored them for an additional 8 and 20 h using established PI staining techniques (45, 51). However, we found no significant changes in cell cycle progression for any of the five cell lines tested (*not shown*). These findings support our model that induction of apoptosis is the primary mechanism by which I3C impairs the proliferation of these cancer cell lines.

### I3C impairs migration and invasive capabilities of cancer cell lines

I3C and BNF have been suggested to interfere with oncogenic signaling pathways that contribute to the establishment of carcinogenic phenotypes (4, 14, 18-20, 30, 52, 53). We sought to investigate whether these compounds could also affect hallmark properties of cancer cell lines, such as migration and invasion. To do so, we tested the migration capability of MDA-MB-231, MCF-7, HCT-8, HEPG2 and HeLa cells in the presence or absence of 4 μM I3C or 10 nM BNF, as determined in our proliferation assays. Cells were grown in 24 well plates and allowed to reach confluency. After serum starvation and incubation with 10 μM AraC to inhibit proliferation, monolayers of three independent biological replicates were scratch-wounded, and cell migration was measured daily (44, 45, 54). Micrographs were taken for several days until control cells closed the wound. While initial migration experiments were performed with 10 μM of I3C, we utilized the lower concentration of 4 μM for these experiments as cells treated with the higher concentration were not able to survive longer than 24-48 h and fully detached from the plates. We observed that I3C had an inhibitory effect on the migratory capabilities of all cell lines, with MCF-7 and HeLa cells being particularly sensitive, and HCT-8 and HEPG2 cell lines showing reduced migration by 35% and 25%, respectively (**Fig. 3**). In contrast, experiments using 10 nM of BNF impaired cell migration only in HepG2 and HeLa cells, with a reduction of 70% and 90%, respectively, compared to controls (**Fig. 3**). Our findings suggest that BNF has a stronger effect than I3C in migration and that it may have a differential effect on the migratory capabilities of different types of breast, ovarian, and hepatocellular carcinoma cells.

**Figure 3.**
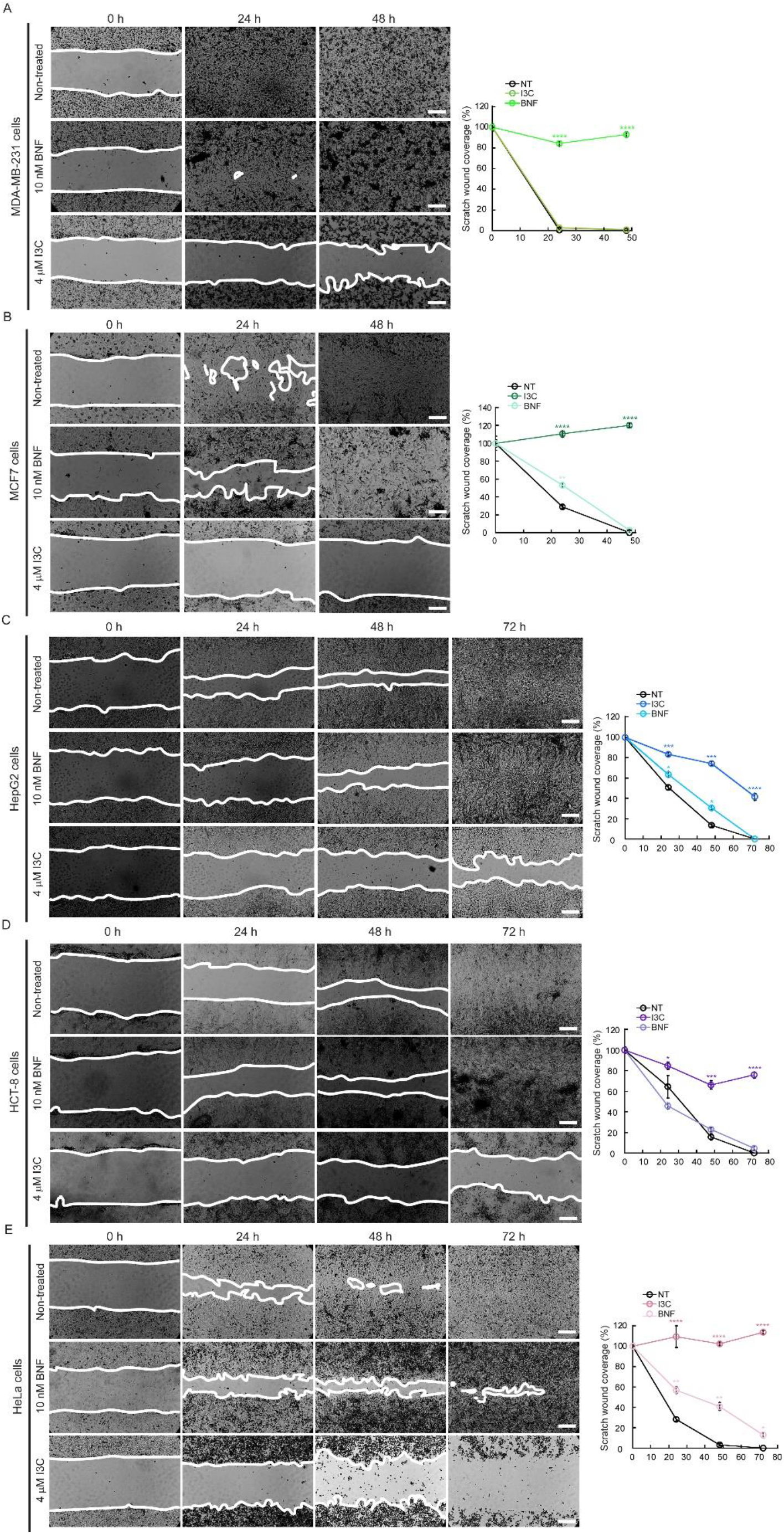
Indole-3-carbinole impairs the directional migration of various cancer cell lines. Representative light microscopy images and quantification of migrated area of wound healing assays of breast cancer cells MDA-MB-231 (**A**) and MCF-7 (**B**), the hepatocellular carcinoma cell lines HepG2 (**C**) and HCT-8 (**D**) and cervical cancer HeLa cells (**E**). Time 0 represents confluent monolayer imaged at the time of performing the wound. The migration of the cells into the wounds was monitored until the monolayers of non-treated cells became fully closed. Images are representative of ten images of three independent biological replicates. Scale bar: 100 μm. For all samples, data are the mean ± SE of three independent biological replicates. *p < 0.05; **p < 0.01; ***p < 0.001; ****p < 0.0001.

In addition to investigating the effect of I3C and BNF on migration, we also examined their impact on the invasive properties of the cancer cell lines using a Matrigel invasion assay (45, 47, 54). We employed the Transwell chamber method and treated three independent biological replicates of each cell line with 10 μM I3C or 10 nM BNF, incubating them for either 24 h (MDA-MD-231, MCF-7, and HeLa cells) or 72 h (HCT-8 and HepG2 cells). The invading cells were then fixed and stained with crystal violet, and non-invading cells were removed from the upper chamber. **Figure 4** illustrates the anti-invasive effect of I3C on all cell lines tested when compared to untreated controls. In breast cancer cell lines, HepG2 and HeLa cells, the effect of I3C was similar to BNF preventing the migration across the Matrigel membrane. However, I3C was less effective than BNF in HCT-8 cells, as there was only a 50% reduction in invasive properties of this cell line when incubated with I3C. These results further support the idea that I3C and BNF can impair the migratory and invasive capabilities of various cancer cell lines, albeit to differing degrees depending on the specific cellular lineage.

**Figure 4.**
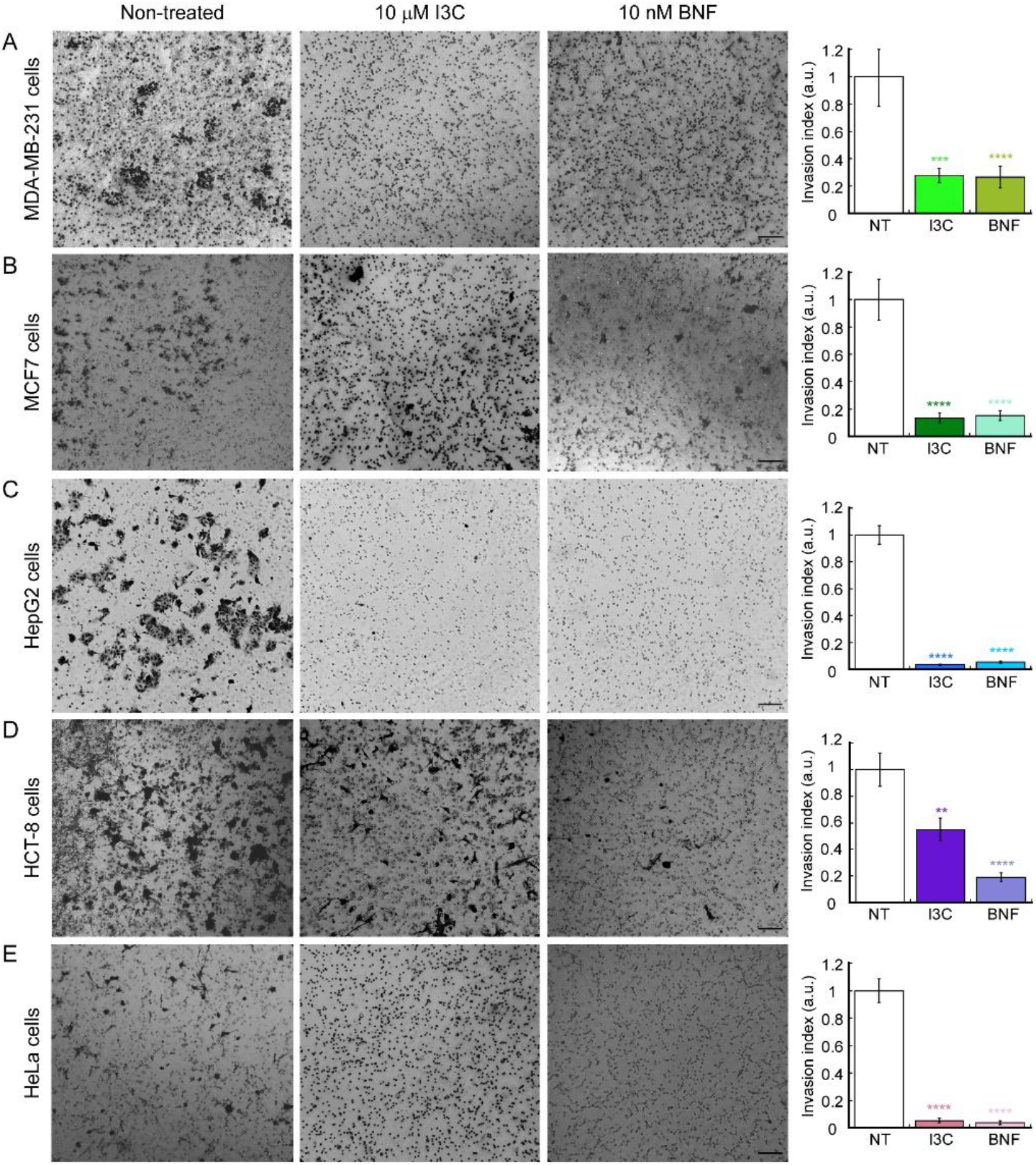
Indole-3-carbinole impairs the invasive properties of various cancer cell lines. Representative light microscopy images (left panels) and quantification (right panels) of Matrigel invasion assay breast cancer cells MDA-MB-231 (**A**) and MCF7 (**B**), the hepatocellular carcinoma cell lines HepG2 (**C**) and HCT-8 (**D**) and cervical cancer HeLa cells (**E**). MDA-MB-231, MCF7 and HeLa cells were incubated for 24 h; HepG2 and HCT-8 cells were cultured for 48 h. The data show the means ± SE of three independent biological replicates imaged and are expressed as the percentage of invading cells compared to the control shown in the plots. Scale bar: 100 μm. For all samples, data are the mean ± SE of three independent biological replicates. **p < 0.01; ***p < 0.001; ****p < 0.0001.

### I3C and BNF impairs mitochondrial potential in cancer cell lines

Cancer cells have a more hyperpolarized membrane potential compared to normal cells, and there seems to be a correlation between the more invasive and aggressive the cancer, with an increased hyperpolarization and decreased metabolism of oxidative phosphorylation (55-58). This phenotype impairs energy production under normal aerobic respiration; however, in most cases cancer cells utilize aerobic glycolysis, via the Warburg effect (59-62). Aerobic glycolysis is a deficient metabolic route to produce ATP as the metabolism of glucose to lactate generates only 2 ATP molecules per glucose utilized. However, it has been proposed that the metabolism of proliferating cancer cells, are adapted to facilitate the incorporation of nutrients into the biological macromolecules essential to generate new cells (59). In this regard, certain signaling pathways related to cell proliferation, and some mutations present in cancer cells might be dedicated to enhance biosynthetic metabolic pathways to better incorporate nutrients into macromolecules important for proliferation instead of energy production (Reviewed by Vander-Heiden, *et al*., 2009 (59)). To assess the effect of I3C and BNF in the metabolic state of the cancer cell lines used here, we performed analyses of mitochondrial membrane potential using the fluorescent dyer TMRE. Proliferating cancer cells were treated with 10 μM I3C and 10 nM BNF for 48 h and then incubated with 200 nM TMRE in the dark for 45 min at 37°C. The cells were then trypsinized and washed three times with PBS, and the fluorescence intensity of TMRE was measured. TMRE is a permeable dye positively-charged that enters and accumulates in active mitochondria, due to the relatively elevated negative charge of this organelle. Loss of mitochondrial function or depolarization results in a reduction of the mitochondrial membrane potential, which can be detected as a reduced accumulation and fluorescence of TMRE dye. The data show that HeLa, MDA-MB-231 and HEPG2 cells that were treated with 10 μM I3C have a significant loss of TMRE fluorescence (**Fig. 5**), which suggest a decreased metabolic state and partially explains the reduced viability observed in these cells (**Fig. 1**). Interestingly, in the case of MCF-7 cells, we detected a significant increase in the TMRE signal, which suggests a different regulatory pathway among ER+ (estrogen receptor positive) and ER- (estrogen receptor negative) cells. BNF treatment had no effect on the mitochondrial potential of any of the cell lines tested here, suggesting that even though both compounds are agonists of AhR, the mechanism of action are different. Studies on the mechanistic links between cell metabolism, regulation of proliferation and associated signaling and gene expression pathways associated to the potential effect of I3C is required to better potential application in treatments against cancer.

**Figure 5.**
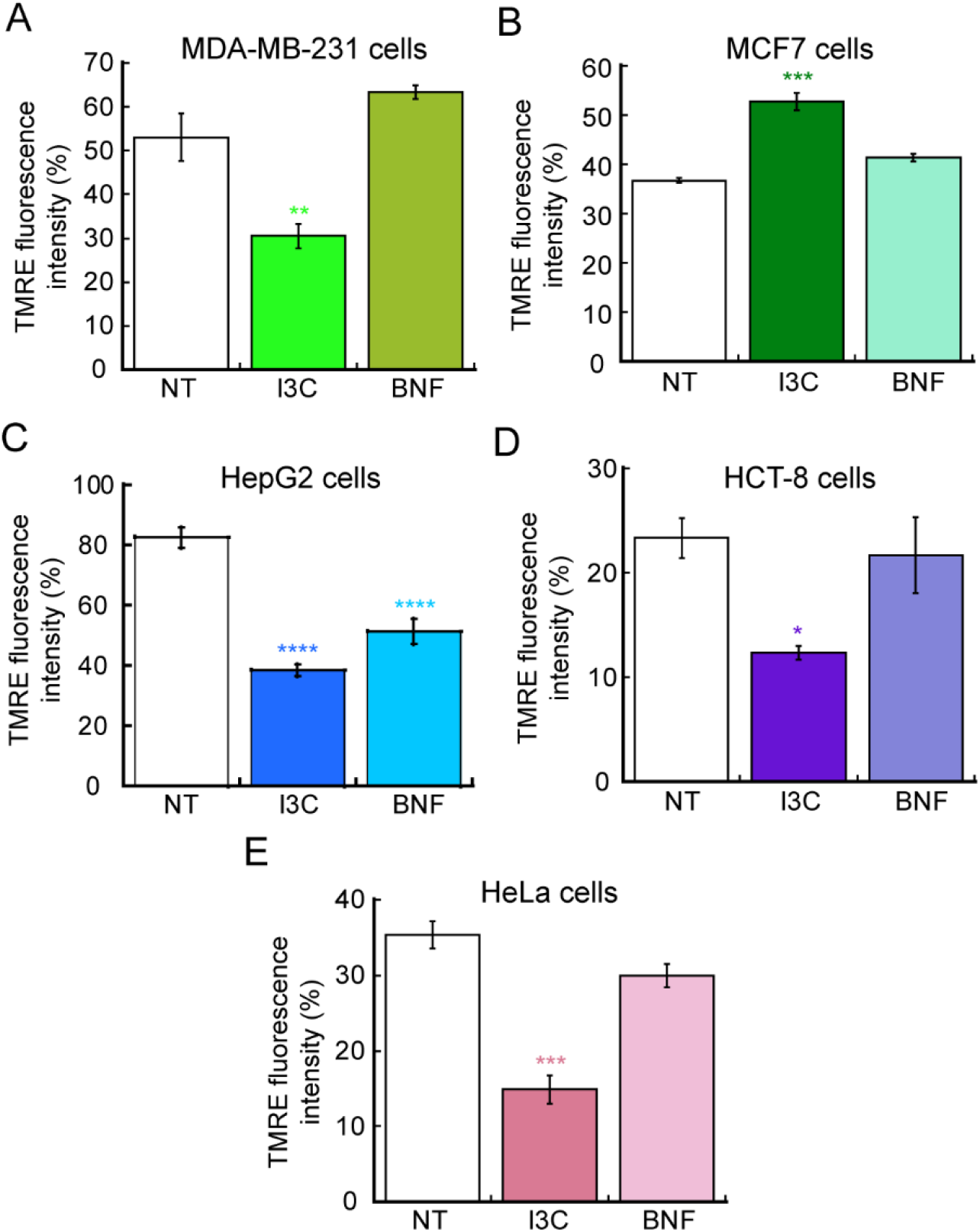
Indole-3-carbinol decreased mitochondrial potential in cancer cell lines. Mitochondrial membrane potential was measured by staining cells with 200 nM TMRE and the percentage of fluorescence intensity of three independent biological replicates was plotted. Breast cancer cells MDA-MB-231 (**A**) and MCF7 (**B**), the hepatocellular carcinoma cell lines HepG2 (**C**) and HCT-8 (**D**) and cervical cancer HeLa cells (**E**) are represented. Data show means ± SE of three independent biological replicates imaged. *p < 0.05; **p < 0.01; ***p < 0.001; ****p < 0.0001 relative to control.

## DISCUSSION

In this study, we observed a positive effect of I3C and BNF on functional hallmarks in various cancer cell lines. Our initial experiments demonstrated that both compounds reduced cellular viability by approximately 50% at concentrations as low as 10 μM for I3C and 5 nM for BNF in ER- and ER+ breast cancer cell lines (MDA-MB-231 and MCF-7), hepatocellular carcinoma models (HCT-8 and HepG2), and cervical cancer (HeLa). While the antiproliferative effects of I3C were also observed in prostate, colon, endometrial, osteosarcoma, and melanoma cancer lines, the IC_50_values for prostate cancer cell lines were higher than those observed in breast and hepatocellular carcinoma cell lines. For instance, studies of I3C in three prostate cancer cell lines (which differ on the levels of p53 expression) such as LNCaP (wild type p53), DU145 (p53 mutant) and PC3 (deficient in p53 gene) were sensitive to this compound at higher doses than what we observed here. The IC_50_ values for LNCaP were 150 μM, 160 μM for DU145, and 285 μM for PC3. In addition, the authors showed that the effect exerted by this drug is partially independent of p53, as in all cases the cells were sensitive to the antiproliferative effects of I3C (12). As a result of activating AhR, I3C and some of its derivatives impairs the proliferation and induce apoptosis of colon, prostate, and endometrial cancer lines as well (9-11, 17, 23, 24, 26, 63, 64). Osteosarcoma and melanoma cells undergo apoptosis when exposed to I3C as well (13, 15). BNF has been shown to also induce anti-proliferative and apoptotic effects in cancer cell lines (8, 29, 30, 52, 65). For instance, BNF has specific effect on ER+ breast cancer cell lines mediated via the AhR and associated downstream PI3K/AKT and MAPK/ERK signaling (66). This report showed an IC_50_ for I3C on growth inhibition of breast cancer cells was 55 μM with a maximal inhibitory effect at 200 μM (66). The variations on concentrations of I3C and BNF in previous studies may be attributed to the purity of the reagents and different exposure times used by the research groups. Moreover, it is known that cancer cell lines can present divergent phenotypes when cultured by different laboratories, indicating the importance of *in vivo* studies to define proper conditions of administration. The observed differential response to I3C and BNF highlights the importance of phenotypical and chemical parameters intrinsic to each cell type, transcriptional expression profiles, signaling pathways, duration of treatment, and dosage. Overall, our study suggests that I3C and BNF have potential as anti-cancer agents and further investigation is warranted.

It is noteworthy that studies have shown that I3C may induce cell cycle arrest and that phytochemical combinatory treatments have an effect over cell cycle progression. For instance, I3C in combination with luteolin induces G1 cell cycle arrest and also inhibit the expression of cyclin-dependent kinase 6 (CDK6) expression in MCF-7 cells (67, 68). However, there is no conclusive data showing a direct effect of I3C in cell cycle. Our gene expression profile suggested that the anti-proliferative effect of the compound was likely due to a larger effect of apoptotic genes, rather than an effect in cell cycle genes. However, it can be hypothesized that a potential indirect effect of I3C on cells cycle arrest might occur under different culture conditions. This idea raises from evidence in breast cancer cells where indole compounds promoted the expression of miRNAs let-7a-e, miR-15a, miR-16, miR-17-5p, miR-19a, and miR-20a and decreased the expression of miR-181a, miR181b, miR-210, miR221 and miR-106a, which ultimately may inactivate various cyclins and impact cell cycle progression (69). However, we found that upon nocodazole arrest and evaluation of various time points, the five cell lines tested here were not responsive to I3C stimulus at the levels of modifying cell cycle behavior. Thus, our model supports an hypothesis where apoptosis is the major contributing factor for decreased growth of cancer cells.

We also investigated the impact of I3C and BNF on the migration and invasion capabilities of five model cancer cell lines. We found that these properties were significantly impaired at lower concentrations than those required to reduce cell growth. Gene expression analyses revealed that I3C primarily induced apoptosis, rather than dysregulation of the cell cycle. Notably, exposure to 5 μM of I3C and 10 nM of BNF resulted in a significant decrease in migration and invasion, with BNF exhibiting greater efficacy at lower doses. Wound healing assays demonstrated that treated cells were unable to migrate and close the wound during the time period in which non-treated control cells healed properly. Consistently, Matrigel invasion assays showed that all cell lines treated with both I3C and BNF were unable to cross the biological substrate and invade the porous membrane, as seen in non-treated control cells. However, the treatment with I3C only reduced migration properties by 50% in HCT-8 cells, suggesting that the mechanism of action of both compounds might differ depending on the cell type. Additionally, we assessed mitochondrial integrity as a proxy measure of metabolic state and found a significant reduction in most cell lines, reflecting impaired metabolism and potentially defective utilization of aerobic glycolysis. Interestingly, opposite mitochondrial behavior was observed in the two breast cancer cell lines tested. While triple-negative MDA-MB-231 cells exhibited a decrease in mitochondrial potential, the ER+ MCF-7 cell line showed an increase of approximately 30% in the intensity of the signal of TMRE. This data is consistent with the fact that metabolically, MCF-7 cells rely on ATP production from oxidative phosphorylation under normal oxygen conditions, and only increase their glycolytic activity under hypoxia (Pasteur type). MDA-MB-231 cells, on the other hand, are considered Warburg type as they utilize aerobic glycolysis for ATP production under both normoxic and hypoxic environments (70, 71). Furthermore, the aryl hydrocarbon receptor nuclear translocator (ARNT) is required for AhR-mediated translocator. ARNT are dimeric partners of several proteins such as the hypoxia inducible factor 1α (HIF1α) (72, 73). Interestingly, BNF down-regulated the expression of ARNT in liver of white sturgeon but an up-regulation in their gill and intestine (74). Therefore, we cannot discard that BNF had an effect over the expression of ARNT resulting in a different effect for this potent AhR ligand than for I3C. This is an important finding, as the differential metabolic response of both cell lines to I3C should be a factor considered when utilizing this compound in different cell lines. Overall, the functional data presented here contributes to the proposal that I3C represents excellent potential natural anti-carcinogenic compound, as it impairs several hallmark phenotypes of cancer.

## CONCLUSION

Our study demonstrated that both I3C and BNF have significant effects on the proliferation, migration, and invasion of various cancer cell lines, with I3C showing effects at a concentration of 10 μM and BNF showing effects at 1 nM. These findings suggest that I3C may serve as a promising complimentary natural therapy for the treatment of cancer. However, further studies using animal models are necessary to confirm its *in vivo* efficacy. I3C not only impairs cell growth and induces apoptosis in various cancer cell models but also affects the mobility and invasive properties of cancer cells, which are crucial for cancer progression and metastasis. The complex molecular interactions of these compounds and their effects on various signaling pathways are important determinants of their efficacy. Therefore, further *in vitro* and *in vivo* studies are needed to determine their potential as primary therapy or co-treatments with existing drugs in cancer patients.

## AUTHOR CONTRIBUTIONS

Conceived and designed the research: T.P.-B., O.D.R.-H. Funding acquisition: A.S.B.-G., T.P.-B., O.D.R.-H. Performed experiments: A.S.B.-G., J.A.C.-C., T.P.-B. Wrote first draft of the manuscript and analyzed data A.S.B.-G., T.P.-B.; Prepared figures and tables A.S.B.-G., T.P.-B.; writing, review and editing, L.I.Q.-G., G.F.-G.; all authors edited and revised and approved the final version of the manuscript.

## ACKNOWLEDGMENTS

The authors are thankful to Ms. Miriam Daniela Zuñiga-Eulogio for her technical support and to Dr. Odette Verdejo-Torres and Monserrat Olea-Flores for their critical comments on this manuscript.

## REFERENCES

1. Organization, W. H. (2021)

2. Observatory, T. G. C. (2020)

3. Dy, G. K., and Adjei, A. A. (2013) Understanding, recognizing, and managing toxicities of targeted anticancer therapies. CA Cancer J Clin 63, 249–279

4. Kim, Y. S., and Milner, J. A. (2005) Targets for indole-3-carbinol in cancer prevention. The Journal of nutritional biochemistry 16, 65–73

5. Fujioka, N., Fritz, V., Upadhyaya, P., Kassie, F., and Hecht, S. S. (2016) Research on cruciferous vegetables, indole-3-carbinol, and cancer prevention: A tribute to Lee W. Wattenberg. Mol Nutr Food Res 60, 1228–1238

6. Tjin Tham Sjin, R. M., Satchi-Fainaro, R., Birsner, A. E., Ramanujam, V. M., Folkman, J., and Javaherian, K. (2005) A 27-amino-acid synthetic peptide corresponding to the NH2-terminal zinc-binding domain of endostatin is responsible for its antitumor activity. Cancer research 65, 3656–3663

7. Shen, B. (2015) A New Golden Age of Natural Products Drug Discovery. Cell 163, 1297–1300

8. Murray, T. J., Yang, X., and Sherr, D. H. (2006) Growth of a human mammary tumor cell line is blocked by galangin, a naturally occurring bioflavonoid, and is accompanied by down-regulation of cyclins D3, E, and A. Breast Cancer Res 8, R17

9. Chinni, S. R., and Sarkar, F. H. (2002) Akt inactivation is a key event in indole-3-carbinol-induced apoptosis in PC-3 cells. Clinical cancer research : an official journal of the American Association for Cancer Research 8, 1228–1236

10. Frydoonfar, H. R., McGrath, D. R., and Spigelman, A. D. (2003) The effect of indole-3-carbinol and sulforaphane on a prostate cancer cell line. ANZ J Surg 73, 154–156

11. Hudson, E. A., Howells, L. M., Gallacher-Horley, B., Fox, L. H., Gescher, A., and Manson, M. M. (2003) Growth-inhibitory effects of the chemopreventive agent indole-3-carbinol are increased in combination with the polyamine putrescine in the SW480 colon tumour cell line. BMC cancer 3, 2

12. Nachshon-Kedmi, M., Yannai, S., Haj, A., and Fares, F. A. (2003) Indole-3-carbinol and 3,3’-diindolylmethane induce apoptosis in human prostate cancer cells. Food Chem Toxicol 41, 745–752

13. Kim, S. Y., Kima, D. S., Jeong, Y. M., Moon, S. I., Kwon, S. B., and Park, K. C. (2011) Indole-3-carbinol and ultraviolet B induce apoptosis of human melanoma cells via down-regulation of MITF. Die Pharmazie 66, 982–987

14. Weng, J. R., Tsai, C. H., Kulp, S. K., and Chen, C. S. (2008) Indole-3-carbinol as a chemopreventive and anti-cancer agent. Cancer Lett 262, 153–163

15. Lee, C. M., Lee, J., Nam, M. J., and Park, S. H. (2018) Indole-3-Carbinol Induces Apoptosis in Human Osteosarcoma MG-63 and U2OS Cells. BioMed research international 2018, 7970618

16. Megna, B. W., Carney, P. R., Nukaya, M., Geiger, P., and Kennedy, G. D. (2016) Indole-3-carbinol induces tumor cell death: function follows form. The Journal of surgical research 204, 47–54

17. Howells, L. M., Gallacher-Horley, B., Houghton, C. E., Manson, M. M., and Hudson, E. A. (2002) Indole-3-carbinol inhibits protein kinase B/Akt and induces apoptosis in the human breast tumor cell line MDA MB468 but not in the nontumorigenic HBL100 line. Mol Cancer Ther 1, 1161–1172

18. Moiseeva, E. P., Fox, L. H., Howells, L. M., Temple, L. A., and Manson, M. M. (2006) Indole-3-carbinol-induced death in cancer cells involves EGFR downregulation and is exacerbated in a 3D environment. Apoptosis 11, 799–812

19. Williams, D. E. (2021) Indoles Derived From Glucobrassicin: Cancer Chemoprevention by Indole-3-Carbinol and 3,3’-Diindolylmethane. Front Nutr 8, 734334

20. Herrmann, S., Seidelin, M., Bisgaard, H. C., and Vang, O. (2002) Indolo[3,2-b]carbazole inhibits gap junctional intercellular communication in rat primary hepatocytes and acts as a potential tumor promoter. Carcinogenesis 23, 1861–1868

21. Nachshon-Kedmi, M., Yannai, S., and Fares, F. A. (2004) Induction of apoptosis in human prostate cancer cell line, PC3, by 3,3’-diindolylmethane through the mitochondrial pathway. British journal of cancer 91, 1358–1363

22. Ge, X., Fares, F. A., and Yannai, S. (1999) Induction of apoptosis in MCF-7 cells by indole-3-carbinol is independent of p53 and bax. Anticancer Res 19, 3199–3203

23. Katdare, M., Osborne, M. P., and Telang, N. T. (1998) Inhibition of aberrant proliferation and induction of apoptosis in pre-neoplastic human mammary epithelial cells by natural phytochemicals. Oncol Rep 5, 311–315

24. Frydoonfar, H. R., McGrath, D. R., and Spigelman, A. D. (2002) Inhibition of proliferation of a colon cancer cell line by indole-3-carbinol. Colorectal Dis 4, 205–207

25. Aggarwal, B. B., and Ichikawa, H. (2005) Molecular targets and anticancer potential of indole-3-carbinol and its derivatives. Cell Cycle 4, 1201–1215

26. Jeon, K. I., Rih, J. K., Kim, H. J., Lee, Y. J., Cho, C. H., Goldberg, I. D., Rosen, E. M., and Bae, I. (2003) Pretreatment of indole-3-carbinol augments TRAIL-induced apoptosis in a prostate cancer cell line, LNCaP. FEBS letters 544, 246–251

27. Bradlow, H. L. (2008) Review. Indole-3-carbinol as a chemoprotective agent in breast and prostate cancer. In Vivo 22, 441–445

28. Nachshon-Kedmi, M., Fares, F. A., and Yannai, S. (2004) Therapeutic activity of 3,3’-diindolylmethane on prostate cancer in an in vivo model. The Prostate 61, 153–160

29. Wang, C., Xu, C. X., Bu, Y., Bottum, K. M., and Tischkau, S. A. (2014) Beta-naphthoflavone (DB06732) mediates estrogen receptor-positive breast cancer cell cycle arrest through AhR-dependent regulation of PI3K/AKT and MAPK/ERK signaling. Carcinogenesis 35, 703–713

30. Anandasadagopan, S. K., Singh, N. M., Raza, H., Bansal, S., Selvaraj, V., Singh, S., Chowdhury, A. R., Leu, N. A., and Avadhani, N. G. (2017) beta-Naphthoflavone-Induced Mitochondrial Respiratory Damage in Cyp1 Knockout Mouse and in Cell Culture Systems: Attenuation by Resveratrol Treatment. Oxid Med Cell Longev 2017, 5213186

31. Hayes, J. D., Kelleher, M. O., and Eggleston, I. M. (2008) The cancer chemopreventive actions of phytochemicals derived from glucosinolates. Eur J Nutr 47 Suppl 2, 73–88

32. Dagnachew Eyachew Amare, T. F. H. B., Patrick P.J. Mulder, Astrid Hamers, Ron L.A.P. Hoogenboom. (2020) Acid condensation products of indole-3-carbinol and their in-vitro (anti)estrogenic, (anti)androgenic and aryl hydrocarbon receptor activities. Arabian Journal of Chemistry, 13, 7199–7211

33. Kim, H. S., and Lee, B. M. J. C. (1997) Inhibition of benzo [a] pyrene-DNA adduct formation by Aloe barbadensis Miller. 18, 771–776

34. Kim, J. Y., Chung, J.-Y., Park, J.-E., Lee, S. G., Kim, Y.-J., Cha, M.-S., Han, M. S., Lee, H.-J., Yoo, Y. H., and Kim, J.-M. J. E. (2007) Benzo [a] pyrene induces apoptosis in RL95-2 human endometrial cancer cells by cytochrome P450 1A1 activation. 148, 5112–5122

35. Vezina, C. M., Lin, T.-M., and Peterson, R. E. J. B. p. (2009) AHR signaling in prostate growth, morphogenesis, and disease. 77, 566–576

36. Brandi, G., Paiardini, M., Cervasi, B., Fiorucci, C., Filippone, P., De Marco, C., Zaffaroni, N., and Magnani, M. (2003) A new indole-3-carbinol tetrameric derivative inhibits cyclin-dependent kinase 6 expression, and induces G1 cell cycle arrest in both estrogen-dependent and estrogen-independent breast cancer cell lines. Cancer research 63, 4028–4036

37. Arellano-Gutierrez, C. V., Quintas-Granados, L. I., Cortes, H., Gonzalez Del Carmen, M., Leyva-Gomez, G., Bustamante-Montes, L. P., Rodriguez-Morales, M., Lopez-Reyes, I., Padilla-Mendoza, J. R., Rodriguez-Paez, L., Figueroa-Gonzalez, G., and Reyes-Hernandez, O. D. (2022) Indole-3-Carbinol, a Phytochemical Aryl Hydrocarbon Receptor-Ligand, Induces the mRNA Overexpression of UBE2L3 and Cell Proliferation Arrest. Curr Issues Mol Biol 44, 2054–2068

38. Wang, C., Xu, C.-X., Bu, Y., Bottum, K. M., and Tischkau, S. A. J. C. (2014) Beta-naphthoflavone (DB06732) mediates estrogen receptor-positive breast cancer cell cycle arrest through AhR-dependent regulation of PI3K/AKT and MAPK/ERK signaling. 35, 703–713

39. Zidi, I., and Balaguer, P. J. M. R. A. (2016) Potential anti-cervical carcinoma drugs with agonist and antagonist AhR/PXR activities. 4

40. Fan, Y., Boivin, G. P., Knudsen, E. S., Nebert, D. W., Xia, Y., and Puga, A. J. C. r. (2010) The aryl hydrocarbon receptor functions as a tumor suppressor of liver carcinogenesis. 70, 212–220

41. Moennikes, O., Loeppen, S., Buchmann, A., Andersson, P., Ittrich, C., Poellinger, L., and Schwarz, M. J. C. r. (2004) A constitutively active dioxin/aryl hydrocarbon receptor promotes hepatocarcinogenesis in mice. 64, 4707–4710

42. Livak, K. J., and Schmittgen, T. D. (2001) Analysis of relative gene expression data using real-time quantitative PCR and the 2(-Delta Delta C(T)) Method. Methods 25, 402–408

43. Verdejo-Torres, O., Flores-Maldonado, C., Padilla-Benavides, T., Campos-Blazquez, J. P., Larre, I., Lara-Lemus, R., Perez Salazar, E., Cereijido, M., and Contreras, R. G. (2019) Ouabain Accelerates Collective Cell Migration Through a cSrc and ERK1/2 Sensitive Metalloproteinase Activity. J Membr Biol

44. Lacombe, M. L., Lamarche, F., De Wever, O., Padilla-Benavides, T., Carlson, A., Khan, I., Huna, A., Vacher, S., Calmel, C., Desbourdes, C., Cottet-Rousselle, C., Hininger-Favier, I., Attia, S., Nawrocki-Raby, B., Raingeaud, J., Machon, C., Guitton, J., Le Gall, M., Clary, G., Broussard, C., Chafey, P., Therond, P., Bernard, D., Fontaine, E., Tokarska-Schlattner, M., Steeg, P., Bieche, I., Schlattner, U., and Boissan, M. (2021) The mitochondrially-localized nucleoside diphosphate kinase D (NME4) is a novel metastasis suppressor. BMC Biol 19, 228

45. Olea-Flores, M., Kan, J., Carlson, A., Syed, S. A., McCann, C., Mondal, V., Szady, C., Ricker, H. M., McQueen, A., Navea, J. G., Caromile, L. A., and Padilla-Benavides, T. (2022) ZIP11 Regulates Nuclear Zinc Homeostasis in HeLa Cells and Is Required for Proliferation and Establishment of the Carcinogenic Phenotype. Frontiers in cell and developmental biology 10, 895433

46. Schindelin, J., Arganda-Carreras, I., Frise, E., Kaynig, V., Longair, M., Pietzsch, T., Preibisch, S., Rueden, C., Saalfeld, S., Schmid, B., Tinevez, J. Y., White, D. J., Hartenstein, V., Eliceiri, K., Tomancak, P., and Cardona, A. (2012) Fiji: an open-source platform for biological-image analysis. Nature methods 9, 676–682

47. Olea-Flores, M., Zuniga-Eulogio, M., Tacuba-Saavedra, A., Bueno-Salgado, M., Sanchez-Carvajal, A., Vargas-Santiago, Y., Mendoza-Catalan, M. A., Perez Salazar, E., Garcia-Hernandez, A., Padilla-Benavides, T., and Navarro-Tito, N. (2019) Leptin Promotes Expression of EMT-Related Transcription Factors and Invasion in a Src and FAK-Dependent Pathway in MCF10A Mammary Epithelial Cells. Cells 8

48. Angireddy, R., Chowdhury, A. R., Zielonka, J., Ruthel, G., Kalyanaraman, B., and Avadhani, N. G. (2020) Alcohol-induced CYP2E1, mitochondrial dynamics and retrograde signaling in human hepatic 3D organoids. Free radical biology & medicine 159, 1–14

49. Chowdhury, A. R., Zielonka, J., Kalyanaraman, B., Hartley, R. C., Murphy, M. P., and Avadhani, N. G. (2020) Mitochondria-targeted paraquat and metformin mediate ROS production to induce multiple pathways of retrograde signaling: A dose-dependent phenomenon. Redox Biol 36, 101606

50. Jump, S. M., Kung, J., Staub, R., Kinseth, M. A., Cram, E. J., Yudina, L. N., Preobrazhenskaya, M. N., Bjeldanes, L. F., and Firestone, G. L. (2008) N-Alkoxy derivatization of indole-3-carbinol increases the efficacy of the G1 cell cycle arrest and of I3C-specific regulation of cell cycle gene transcription and activity in human breast cancer cells. Biochem Pharmacol 75, 713–724

51. Padilla-Benavides, T., Olea-Flores, M., Nshanji, Y., Maung, M.T., Syed, S.A. and Imbalzano, A.N. (2022) Differential requirements for different subfamilies of the mammalian SWI/SNF chromatin remodeling enzymes in myoblast cell cycle progression and expression of the Pax7 regulator. Biochimica et Biophysica Acta (BBA) - Gene Regulatory Mechanisms

52. Prochaska, H. J., and Talalay, P. (1988) Regulatory mechanisms of monofunctional and bifunctional anticarcinogenic enzyme inducers in murine liver. Cancer research 48, 4776–4782

53. Takahashi, N., Dashwood, R. H., Bjeldanes, L. F., Williams, D. E., and Bailey, G. S. (1995) Mechanisms of indole-3-carbinol (I3C) anticarcinogenesis: inhibition of aflatoxin B1-DNA adduction and mutagenesis by I3C acid condensation products. Food Chem Toxicol 33, 851–857

54. Nagle, I., Richert, A., Quinteros, M., Janel, S., Buysschaert, E., Luciani, N., Debost, H., Thevenet, V., Wilhelm, C., Prunier, C., Lafont, F., Padilla-Benavides, T., Boissan, M., and Reffay, M. (2022) Surface tension of model tissues during malignant transformation and epithelial-mesenchymal transition. Frontiers in cell and developmental biology 10, 926322

55. Fantin, V. R., Berardi, M. J., Scorrano, L., Korsmeyer, S. J., and Leder, P. (2002) A novel mitochondriotoxic small molecule that selectively inhibits tumor cell growth. Cancer Cell 2, 29–42

56. Nguyen, C., and Pandey, S. (2019) Exploiting Mitochondrial Vulnerabilities to Trigger Apoptosis Selectively in Cancer Cells. Cancers (Basel) 11

57. Bonnet, S., Archer, S. L., Allalunis-Turner, J., Haromy, A., Beaulieu, C., Thompson, R., Lee, C. T., Lopaschuk, G. D., Puttagunta, L., Bonnet, S., Harry, G., Hashimoto, K., Porter, C. J., Andrade, M. A., Thebaud, B., and Michelakis, E. D. (2007) A mitochondria-K+ channel axis is suppressed in cancer and its normalization promotes apoptosis and inhibits cancer growth. Cancer Cell 11, 37–51

58. Badrinath, N., and Yoo, S. Y. (2018) Mitochondria in cancer: in the aspects of tumorigenesis and targeted therapy. Carcinogenesis 39, 1419–1430

59. Vander Heiden, M. G., Cantley, L. C., and Thompson, C. B. (2009) Understanding the Warburg effect: the metabolic requirements of cell proliferation. Science 324, 1029–1033

60. DeBerardinis, R. J., and Chandel, N. S. (2020) We need to talk about the Warburg effect. Nat Metab 2, 127–129

61. Liu, C., Jin, Y., and Fan, Z. (2021) The Mechanism of Warburg Effect-Induced Chemoresistance in Cancer. Frontiers in oncology 11, 698023

62. Warburg, O. (1956) On the origin of cancer cells. Science 123, 309–314

63. Hong, C., Firestone, G. L., and Bjeldanes, L. F. (2002) Bcl-2 family-mediated apoptotic effects of 3,3’-diindolylmethane (DIM) in human breast cancer cells. Biochem Pharmacol 63, 1085–1097

64. Jinno, H., Steiner, M. G., Mehta, R. G., Osborne, M. P., and Telang, N. T. (1999) Inhibition of aberrant proliferation and induction of apoptosis in HER-2/neu oncogene transformed human mammary epithelial cells by N-(4-hydroxyphenyl)retinamide. Carcinogenesis 20, 229–236

65. Fernandes, C., Horn, A., Jr., Lopes, B. F., Bull, E. S., Azeredo, N. F. B., Kanashiro, M. M., Borges, F. V., Bortoluzzi, A. J., Szpoganicz, B., Pires, A. B., Franco, R. W. A., Almeida, J. C. A., Maciel, L. L. F., Resende, J., and Schenk, G. (2015) Induction of apoptosis in leukemia cell lines by new copper(II) complexes containing naphthyl groups via interaction with death receptors. Journal of inorganic biochemistry 153, 68–87

66. Fares, F. A., Ge, X., Yannai, S., and Rennert, G. (1998) Dietary indole derivatives induce apoptosis in human breast cancer cells. Advances in experimental medicine and biology 451, 153–157

67. Wang, X., Zhang, L., Dai, Q., Si, H., Zhang, L., Eltom, S. E., and Si, H. (2021) Combined Luteolin and Indole-3-Carbinol Synergistically Constrains ERalpha-Positive Breast Cancer by Dual Inhibiting Estrogen Receptor Alpha and Cyclin-Dependent Kinase 4/6 Pathway in Cultured Cells and Xenograft Mice. Cancers (Basel) 13

68. Cram, E. J., Liu, B. D., Bjeldanes, L. F., and Firestone, G. L. (2001) Indole-3-carbinol inhibits CDK6 expression in human MCF-7 breast cancer cells by disrupting Sp1 transcription factor interactions with a composite element in the CDK6 gene promoter. J Biol Chem 276, 22332–22340

69. El-Daly, S. M., Gamal-Eldeen, A. M., Gouhar, S. A., Abo-Elfadl, M. T., and El-Saeed, G. (2020) Modulatory Effect of Indoles on the Expression of miRNAs Regulating G1/S Cell Cycle Phase in Breast Cancer Cells. Applied biochemistry and biotechnology 192, 1208–1223

70. Sakamoto, T., Niiya, D., and Seiki, M. (2011) Targeting the Warburg effect that arises in tumor cells expressing membrane type-1 matrix metalloproteinase. J Biol Chem 286, 14691–14704

71. Gatenby, R. A., and Gillies, R. J. (2004) Why do cancers have high aerobic glycolysis? Nat Rev Cancer 4, 891–899

72. Hoffman, E. C., Reyes, H., Chu, F.-F., Sander, F., Conley, L. H., Brooks, B. A., and Hankinson, O. J. S. (1991) Cloning of a factor required for activity of the Ah (dioxin) receptor. 252, 954–958

73. Gonzalez, F. J., Xie, C., and Jiang, C. (2019) The role of hypoxia-inducible factors in metabolic diseases. Nature Reviews Endocrinology 15, 21–32

74. Doering, J. A., Beitel, S. C., Patterson, S., Eisner, B. K., Giesy, J. P., Hecker, M., and Wiseman, S. (2020) Aryl hydrocarbon receptor nuclear translocators (ARNT1, ARNT2, and ARNT3) of white sturgeon (Acipenser transmontanus): Sequences, tissue-specific expressions, and response to β-naphthoflavone. Comparative Biochemistry and Physiology Part C: Toxicology & Pharmacology 231, 108726

